# Prolonged deficit of gamma oscillations in the peri-infarct cortex of mice after stroke

**DOI:** 10.1101/2020.03.05.978593

**Authors:** M Hazime, M Alasoadura, R Lamtahri, P Quilichini, J Leprince, D Vaudry, J Chuquet

## Abstract

Days and weeks after an ischemic stroke, the peri-infarct area adjacent to the necrotic tissue exhibits very intense synaptic reorganization aimed at regaining lost functions. In order to enhance functional recovery, it is important to understand the mechanisms supporting neural repair and neuroplasticity in the cortex surrounding the lesion. Brain oscillations of the local field potential (LFP) are rhythmic fluctuations of neuronal excitability aimed at synchronizing neuronal activity to organize information processing and plasticity. Although the oscillatory activity of the brain has been probed after stroke in both animals and humans using electroencephalography (EEG), the latter is ineffective to precisely map the oscillatory changes in the peri-infarct zone where synaptic plasticity potential is high. Here, we worked on the hypothesis that brain oscillatory system is altered in the surviving peri-infarct cortex, which may slow down functional repair and reduce the capacity to recovery. In order to document the relevance of this hypothesis, oscillatory power was measured at various distances from the necrotic core at 7 and 21 days after a permanent cortical ischemia induced in mice. Delta and theta oscillations remained at a normal power in the peri-infarct cortex, in contrast to gamma oscillations that displayed a rapid decrease, the closer we get to the lesion core. A broadband increase of power was also observed in the homotopic contralateral sites. Thus, the proximal peri-infarct cortex could become a target of therapeutic interventions aimed at correcting the oscillatory regimen. These results argue for the usefulness of therapeutic intervention aimed at boosting gamma oscillations in order to improve post-stroke functional recovery.

## Introduction

Stroke remains the most frequent cause of disability stemming from the lack of efficient therapy to increase the magnitude of functional recovery. In the chronic stage following a focal brain ischemia, neuronal plasticity possibly leads to a regain of lost functions. The wiring of new circuits uses both established and *de novo* cellular materials, from spinogenesis to axonal sprouting (Brown et al., 2009; Carmichael et al., 2001; Florence et al., 1998; Murphy and Corbett, 2009; Nudo and Milliken, 1996; Nudo et al., 1996; Stroemer et al., 1998; Ueno et al., 2012). The reorganization of neuronal networks potentially occurs in any structure involved in the relearning of a behavior. Intense plastic changes take place especially in the cortex adjacent to the ischemic lesion, the so-called peri-infarct cortex, where neurons are well alive but neural circuits are partially damaged (Brown et al., 2007; Murphy and Corbett, 2009; Stroemer et al., 1995). There is an increasing number of studies showing how the somatosensory-motor cortical activities of the affected limbs are progressively transferred and reorganized in the peri-infarct area (Brown et al., 2009; Carmichael, 2012; Dijkhuizen et al., 1999; Sharma and Cohen, 2012; Tombari et al., 2004; Wang et al., 2009). Progress from basic research also suggests that cellular and molecular mechanisms aimed at assembling neurons after stroke are similar to those identified during learning and memory formation in physiological context (*e.g*. long-term change of synaptic strength) (Chervyakov et al., 2015; Cooke and Bliss, 2006; Yu et al., 2019). However, the peri-infarct cortex is under the influence of various pathophysiological processes such as chronic inflammation and glial reactivity. It is not yet clear how and to what extent these micro-environmental conditions interfere with neuronal reconnection (Skaper et al., 2018; Sochocka et al., 2017). For instance, neuronal excitability, a determining factor for neuronal communication and synaptic plasticity, is reduced in the peri-infarct area, which has been demonstrated to limit functional recovery (Clarkson et al., 2010; Carmichael, 2012). Thus, a therapeutic goal to optimize recovery after acute brain damage is to normalize the conditions of neuronal communication.

Neural oscillations are rhythmic fluctuations of the brain electrical field potential (including membrane potential of neurons) that can be measured with extracellular electrodes placed into the cortex (local field potential, LFP) or with external electrodes placed over the scalp (electroencephalography, EEG). Several discrete frequency bands generated by different oscillators have been identified, from slow rhythms (*e.g*. delta band, 1-3 Hz) to fast rhythms (*e.g*. gamma band, 30-80 Hz). Each rhythm is more or less expressed depending on the brain state (*i.e*. wake *vs*. sleep) and the ongoing sensory, motor or cognitive activity. The fundamental role of these rhythmic fluctuations of neuronal excitability is to regulate action potentials trafficking by synchronizing the firing of multiple neuronal assemblies distributed across brain regions (Buzsáki 2007). Interestingly, an increasing number of studies show the potential importance of oscillations in neuronal plasticity (Womelsdorf & Hoffman, 2018; Zarnadze et al., 2016). For instance, reduction of theta (3-7 Hz) and gamma (30-80 Hz) activity has been correlated with decreased long term potentiation (LTP) (Huerta & Lisman, 1993; Kalemaki et al., 2018; Kalweit et al., 2017; Stephan et al., 2009; Wespatat et al., 2004; Huerta and Lisman, 1993; Wespatat et al., 2004; Stephan et al., 2009; Kalweit et al., 2017; Kalemaki et al., 2018). Indeed, theta and gamma wave-bands are precisely tuned to dictate a timing of collective neuronal discharges that match the temporality of synaptic plasticity mechanisms, such as LTP or spike timing dependent plasticity (STDP) (Bi and Poo, 1998; Buzsáki and Draguhn, 2004; Engel et al., 1992; Harris et al., 2005, 2003; König et al., 1996; Magee and Johnston, 1997; Wespatat et al., 2004; Nishiyama et al., 2010). Logically, the modification of network connections allowing functional recovery after a stroke requires that homeostatic mechanisms such as the maintenance of neuronal oscillatory activity stick to their operating range (Buzsáki, 2006; Llinás et al., 2005; Schnitzler & Gross, 2005). Several tools are tested with the aim to precisely tune the oscillatory spectrum of the pathological brain by the mean of pharmacologic, optogenetic, transcranial magnetic, direct current or rhythmic sensory stimulation methods (Guerra et al., 2018; Lozano-Soldevilla, 2018). Therefore, it is important to document the oscillatory conditions of the repairing cortex after stroke. In human, measuring the rhythmic activity of the post-stroke cortex is not new (Carino-Escobar et al., 2019; Giaquinto et al., 1994; Zhang et al., 2013) and in most cases the purpose was to find an early biomarker able to predict the quality of recovery (Laaksonen et al., 2013; Nicolo et al., 2015; Pfurtscheller, 1981; Tecchio et al., 2005; Van Huffelen et al., 1984; Zhang et al., 2013; Rabiller et al., 2015;). But these studies were carried out with EEG or magnetoencephalography (MEG), whose spatial resolution is too low to probe the ~2 mm peri-infarct boundary extending from the edge of the core (Ohab et al., 2006; Brown et al., 2007). Therefore, little is known about the spatial extent of neural oscillatory changes around the necrotic neuronal mass. The aim of the present work was thus to describe how the oscillatory activity of the peri-infarct cortex evolves, one and three weeks after a focal cortical ischemia in mice.

## Material and Methods

### Study approval

Experiments, approved by the Ethics Committee for Animal Research of Normandy (CENOMEXA) and the French ministry of higher education, research and innovation, were conducted by authorized investigators in accordance with the recommendations of the European Community Council Directive 2010/63/UE of September 22.2010 on the protection of animals used for scientific purposes.

### Stroke model

For these studies, 8-12-week-old (20-25g) male C57BL/6J mice were purchased from Janvier Laboratories. Anesthesia was induced with 5% isoflurane in an induction chamber. Following the loss of righting reflex, the mouse was rapidly transferred to a nose cone mask, and maintained with isoflurane (2-2.5%) delivered at 1 L/min in oxygen enriched air and 0.15 mg/kg of buprenorphine (Par Sterile Products) was injected i.p. Spontaneous breathing frequency was kept over 0.5 Hz and body temperature was strictly maintained at 37 ± 0.2°C throughout the surgery. Focal ischemia was induced by the permanent occlusion of the left middle cerebral artery as reported previously (Nawashiro et al., 1997; Welsh et al., 1987). Briefly, a skin incision was made between the orbit and ear. Under an operating microscope, an incision was made, dividing the temporal muscle, and the left lateral aspect of the skull was exposed by reflecting the temporal muscle and surrounding soft tissue. The distal course of the middle cerebral artery was then visible through the translucent skull. A small burr-hole craniectomy was performed with a dental drill. The left middle cerebral artery was electrocoagulated by bipolar diathermy. The muscle and soft tissue were replaced, and the incision was sutured. In this model, ischemia is restricted to the neocortex. An analgesic (bupreorphine) was administered on the day of the surgery (0.015 mg/kg, s.c.) and on subsequent 2 days.

### *In vivo* electrophysiology

Seven or 21 days after the occlusion of the middle cerebral artery, anesthesia was induced with 5% isoflurane in an induction chamber. Following the loss of righting reflex, the mouse was rapidly transferred to a nose cone mask and maintained with isoflurane (2-2.5%) delivered at 1 L/min in oxygen enriched air. A calibrated isoflurane dispenser with consistent and constant oxygen pressure was used to ensure reliable and reproducible delivery of specific isoflurane concentrations. Animal was placed in a stereotaxic frame (Kopft Instrument), and 0.15 mg/kg of buprenorphine (Par Sterile Products) was injected i.p. and the eyes were protected with ophthalmic gel. Breathing frequency was maintained over 0.5 Hz and body temperature was strictly maintained at 37 ± 0.2°C throughout the surgery using a feedback-controlled heating pad. The head was shaved and disinfected with iodine, the cranium was exposed, and 3 small holes were drilled in the parietal bone and the dura-matter was incised with care to leave the cortex intact. Three Teflon-coated silver wire electrodes, 100 μm in diameter, were lowered into the cortex until layer 4. Two electrodes were placed in the ischemic ipsilateral hemisphere at varying distances (<2.5mm) from the putative lesion border. The third electrode was placed in the contralateral hemisphere in homotopic position with the distal ipsilateral electrode. The 3 electrodes were secured to the skull with some dental cement (Paladur, DentalMedical). Two other electrodes were implanted in the cerebellum for reference and ground readings. After the surgical procedure, isoflurane was reduced to 1.1 ± 0.1% for a resting period of 45-60 min followed by the recording period. Fifteen minutes of recording were performed twice with each electrode resulting in 6 different readings overall. At the end, anesthesia was elevated to 5% for a 5 min period followed by an intracardiac injection of saturated KCl solution inducing heart arrest.

### Data acquisition and analysis

All *in vivo* recordings were done in a Faraday chamber and used an amplifier PowerLab 8/35 (AD-Instrument). Raw data were acquired with Labchart software (AD Instrument). The extracellular signals were amplified (10000×), bandpass filtered (0.1 Hz to 100 Hz) for the local field potential (LFP) and acquired continuously at 4 kHz. In addition, a notch filter was added to eliminate the 50 Hz band noise. For spectral analysis of oscillatory patterns (epoch of 15 min), a modified version of the multitaper FFT MATLAB package by Mitra and Pesaran (1999) was used: FFT window size of 4 s, three to five tapers, frequency bins = 0.15 Hz, no overlap between successive windows, time bandwidth = 3 (Mitra and Pesaran, 1999; Quilichini et al., 2010).

### Measurement of Infarct-electrodes distance

After the recording, under deep anesthesia, mice were killed, and 1 μL of black ink was slowly injected using a micropipette into the site of each ipsilateral electrode. The brains were removed rapidly and frozen in cooled isopentane at −65°C for 10 seconds. Whole brains were cut in 20-μm-thick sections with a cryostat (Leica CM3050), and 1/5 serial sections were collected on a glass slide and stained by thionine. Thionine stained slides were scanned at a high resolution and imported into ImageJ. The Euclidian distance between ink spots and the closer lesion border was calculated.

### Immunofluorescent labeling of GFAP

Reactive astrogliosis was assessed by immunofluorescent labeling for GFAP at 7 and 21 days post stroke. Brains were sectioned and every 5^th^ section (20 μm thick) was collected and stored in cryopreservation solution until used. Sections were rinsed in buffered saline and transferred to 1% BSA in tris-buffered saline (TBS) for 10 min at room temperature, followed by a second TBS rinse. The sections were blocked for 30 min in TBS containing 2% donkey serum with 0.3% Triton X-100 before being incubated in TBS with 2% donkey serum and 0.3% triton X-100 containing the primary antibody rabbit GFAP, 1:600 (ref Z033401-2, Agilent Dako) for 24 hours at 4°C. Ensuing incubation with primary antibody, sections were washed three times in TBS followed by incubation for 2 hours at room temperature in TBS with 2% normal donkey serum and 0.3% triton X-100 containing the fluorescent secondary antibody (Donkey anti rabbit Alexa-488, 1:300, A-21206, Invitrogen) and finally by the nuclear counter stain DAPI (1:1000, Sigma-Aldrich) in TBS for 5 min at room temperature. Images were taken with a Leica SP8 microscope, where two sections from each animal (n = 3) were included in the analysis.

### Statistics

Data are presented as mean ± SEM. All statistics were performed using Prism software (Graphpad). A paired Student’s *t*-test was used for pairwise comparisons.

## Results

Cerebral ischemia was induced in mice using a variation of the well-characterized Tamura model (Tamura et al., 1981) consisting in a permanent occlusion of a distal trunk of the middle cerebral artery. This procedure generated a reproducible focal lesion (8.5 ± 0.8 mm^3^ at 48 h, n = 20 mice) representing about 12% of the volume of the entire ipsilateral cortex. Sham operated animals didn’t present any lesion. In order to visualize the extent of the pathophysiological alteration of the surviving cortex adjacent to the necrotic core, the cellular reactivity of astrocytes, an endogenous biomarker for brain diseases, was investigated. Upregulation of GFAP is generally considered as a reliable maker for reactive astrocytes (Escartin et al., 2019). The immunostaining of GFAP showed a gradual decrease in intensity as we move away from the stroke border (Figure 1). At day 7 post-stroke, the up-expression of GFAP was most intense in a 315 ± 12 μm (n = 6 mice) zone surrounding the necrotic core, where a long leading process of each astrocyte was extended in the direction of the infarct (Figure 1A). Further away, a second peripheral area of 1552 ± 52 μm was defined by intense GFAP staining with protoplasmic shape astrocytes without preferred orientation of their processes (Figure 1A). Beyond, GFAP expression was no more different from its expression in the contralateral cortex (Figure 1Aa). At day 21, the glial scar was narrower (163 ± 3 μm) but overall, the glially-defined peri-infarct area was more extended (2320 ± 40 μm) (Figure 1B).

**Figure 1.**
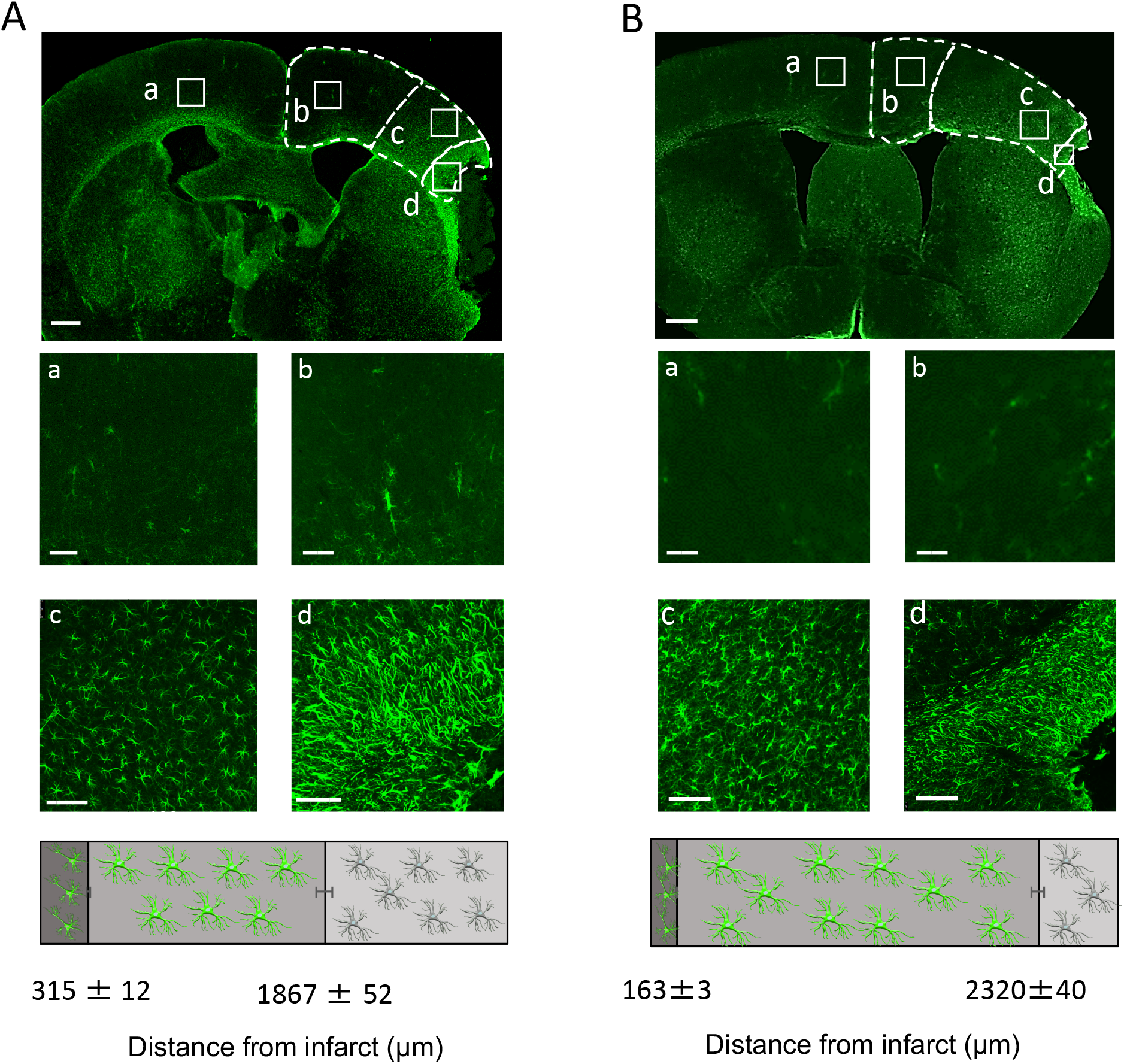
Astroglial identification of the peri-infarct cortex. GFAP immunostaining of a coronal section of the mouse brain 7 days (**A**) and 21 days (**B**) after MCAO. Four regions of interest were identified: **a**. the contralateral cortex where GFAP immunoreactivity is low; **b**. the distal ipsilateral cortex where the level of GFAP expression is as low as in the contralateral cortex; **c**. the proximal ipsilateral cortex where astrocytes are strongly GFAP-positive; **d**. the border of the infarction, the glial scar, where astrocytes express high level of GFAP with a long leading process oriented toward the lesion (or the cavity). Hence, 3 distinct zones were define, the extent of which is reported in the horizontal stacked bar down to panel A and B (average Euclidian length of each zone ± SEM; n = 3 mice per time point).

To test the hypothesis of an alteration of neural oscillatory activity around the cortical infarction, 3 electrodes were implanted in the cortex of the mouse brain, 7 or 21 days after MCAO. Two electrodes (E1 and E2) were placed in cortical layer 4 at various distances (< 4 mm) from the expected location of the lesion border (Figure 2A). The third electrode (E3) was placed in the layer 4 of the contralateral hemisphere, in mirror position with the nearest electrode from the lesion. In sham animals, two electrodes were place in mirror position, in the layer 4 of the barrel field cortex. After 45-60 min of rest under isoflurane anesthesia (1.1 ± 0.1%) and during which body temperature was strictly kept at 37 ± 0.2°C, each recording was done twice during 15 min at 7 days or 21 days. In such condition, the whitened power spectrum of LFP activity shows 3 distinct peaks at 1.8 ± 0.8 Hz, 4.5 ± 0.9 Hz and 36.4 ± 1.1 Hz, corresponding respectively to the delta (1 – 3 Hz), the theta (4 – 7 Hz) and the gamma (30 – 50 Hz) waveband (Figure 2B). The other wavebands (alpha, beta, high gamma) were considered inactive. Sham operated animals didn’t show any LFP spectrum power difference between the ipsilateral and the contralateral hemisphere (*P* > 0.05 for the three wavebands at both 7 and 21 days post-stroke, n = 4 animals, Figure 2B). Irrespective to their distance from the infarct border, averaging the power spectrum recorded by all ipsilateral electrodes showed a slight depression of the broadband cortical oscillatory power at day 7 post-stroke, compared with sham animal, albeit not significant (*P* > 0.05 for the 3 wavebands studied; n = 8 ipsilateral electrodes from 4 mice, Figure 2B). In contrast, the contralateral hemisphere showed a global increase in power, relative to sham animal. As a result, at 7 days, there was a significant difference of the theta and the gamma waveband power between the contralateral and the ipsilateral (66.8 ± 0.5 dB vs. 63.3 ± 1.8 dB for theta, *P* < 0.01;

72.5 ± 1.1 vs. 67.3 ± 1.5 for gamma, *P* < 0.05; n = 8 ipsilateral electrodes from 4 mice Figure 2B). At day 21 post-stroke, the ipsilateral power spectrum (irrespective to electrodes position) moved toward normalization relative to sham (Figure 2C), while the LFP power of the contralateral cortex remained significantly increased for the delta (P < 0.01), the theta (P < 0.05) and the gamma wavebands relative to the ipsilateral LFP power (P < 0.05; n = 5 contralateral electrodes and n = 10 ipsilateral electrodes from 5 mice, Figure 2C and D).

**Figure 2.**
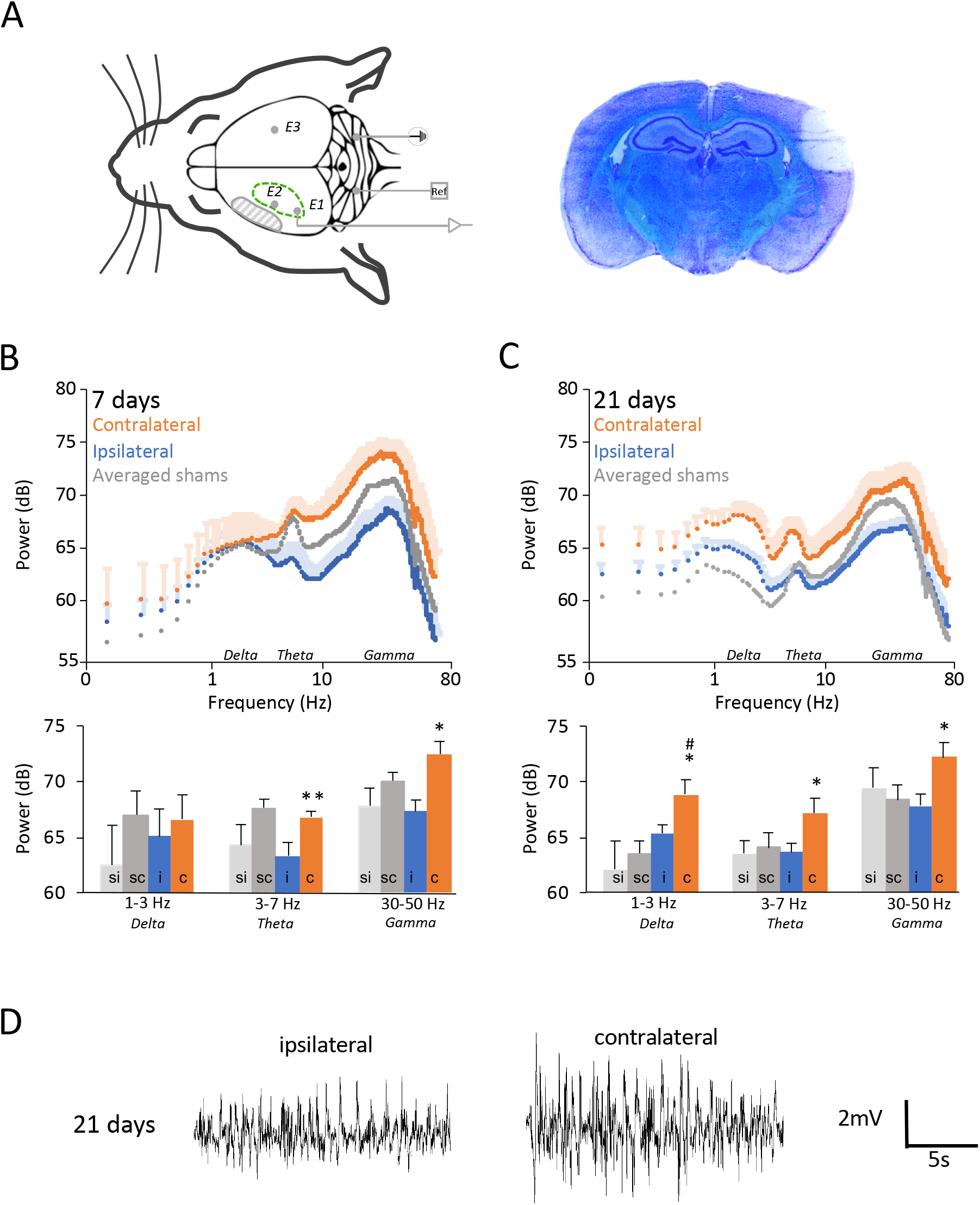
Spectral analysis of the LFP activity of the peri-infarct cortex. **A**. Left: experimental arrangement. Three electrodes were implanted for sequential recordings of 15 min in the ipsilateral cortex (E1 and E2) and in the contralateral cortex (E3); R: reference electrode. Right: a representative ischemic lesion 48h after the MCAO procedure. Whitened power spectrum (1-80 Hz) obtains at 7 days (**B**) and 21 days (**C**) in sham and ischemic animal. Note the three distinct peaks corresponding to the frequencies of delta, theta and gamma waveband. To clarify the graphical representation, the ipsi- and contra-lateral power spectrums of sham animals where averaged and presented without error bars. The power quantification of each waveband is reported in the respective histogram below. Comparisons between hemispheres (ipsi-vs. contra-lateral) and between sham and stroke animals are reported by (*) and (^#^) respectively when significant (* or ^#^ *P* < 0.05; ** *P* < 0.01; 7 days: n = 8 recordings from 4 ischemic animals at 7 days post-stroke and n = 10 recordings from 5 animals at 21 days post-stroke. Sham animals: n = 4 animal per time point). **D**. Representative raw trace (1-80Hz band pass filtered) of the LFP activity of the ipsilateral and contralateral cortex recorded 21 days after MCAO.

To investigate the spatial relation between the oscillatory activity and the extent of the glially defined peri-infarct area, the scar border-to-electrodes distances were measured precisely by histological analysis, using ink labeling as a marker of the electrode sites. No significant correlation between the ipsilateral LFP power and the distance from the infarct border was found for the delta and the theta waveband at 7 or 21 days post-stroke (7 days: *P* > 0.05, n = 8 electrodes from 4 mice; 21 days: *P* > 0.05, n = 10 electrodes from 5 mice; Figure 3A and B). However, for the gamma waveband (30-50 Hz), there was a significant positive relationship between LFP power and distance to the infarct border at both 7 and 21 days post-stroke (7 days: r(6) = 0.912; *P* < 0.05, n = 8 electrodes from 4 mice; 21 days: r(8) = 0.752; *P* < 0.05, n = 10 electrodes from 5 mice; Figure 3A and B). The relationship was best described by a power function which equation was used to build a visual model showing the spatial extent of the gamma depression relative to the average location and size of the ischemic lesion (Figure 4).

**Figure 3.**
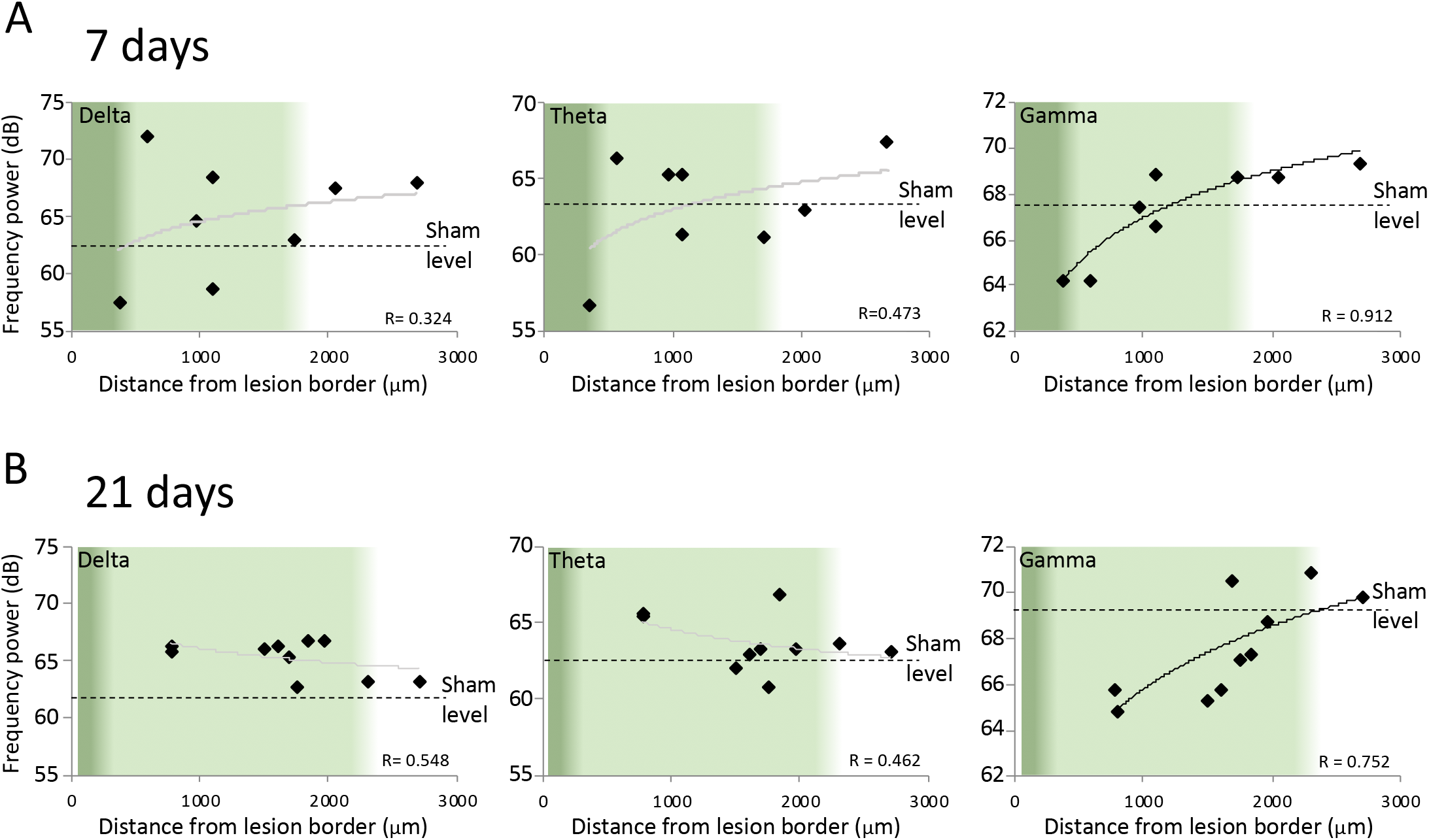
Analysis of LFP power change relative to the distance from the cortical infarct. At each time point, (**A:** 7 days; **B:** 21 days) LFP power and the distance between peri-infarct electrodes and the lesion border where plotted to test the hypothesis of a correlation. No correlation where found for delta and theta. However, the power of gamma oscillation (30-50 Hz) showed a strong correlation with the distance from the infarct border. The green background indicates the limits of the astroglial scar (dark green) and the peri-infarct zone (light green) as reported in figure 1.

**Figure 4.**
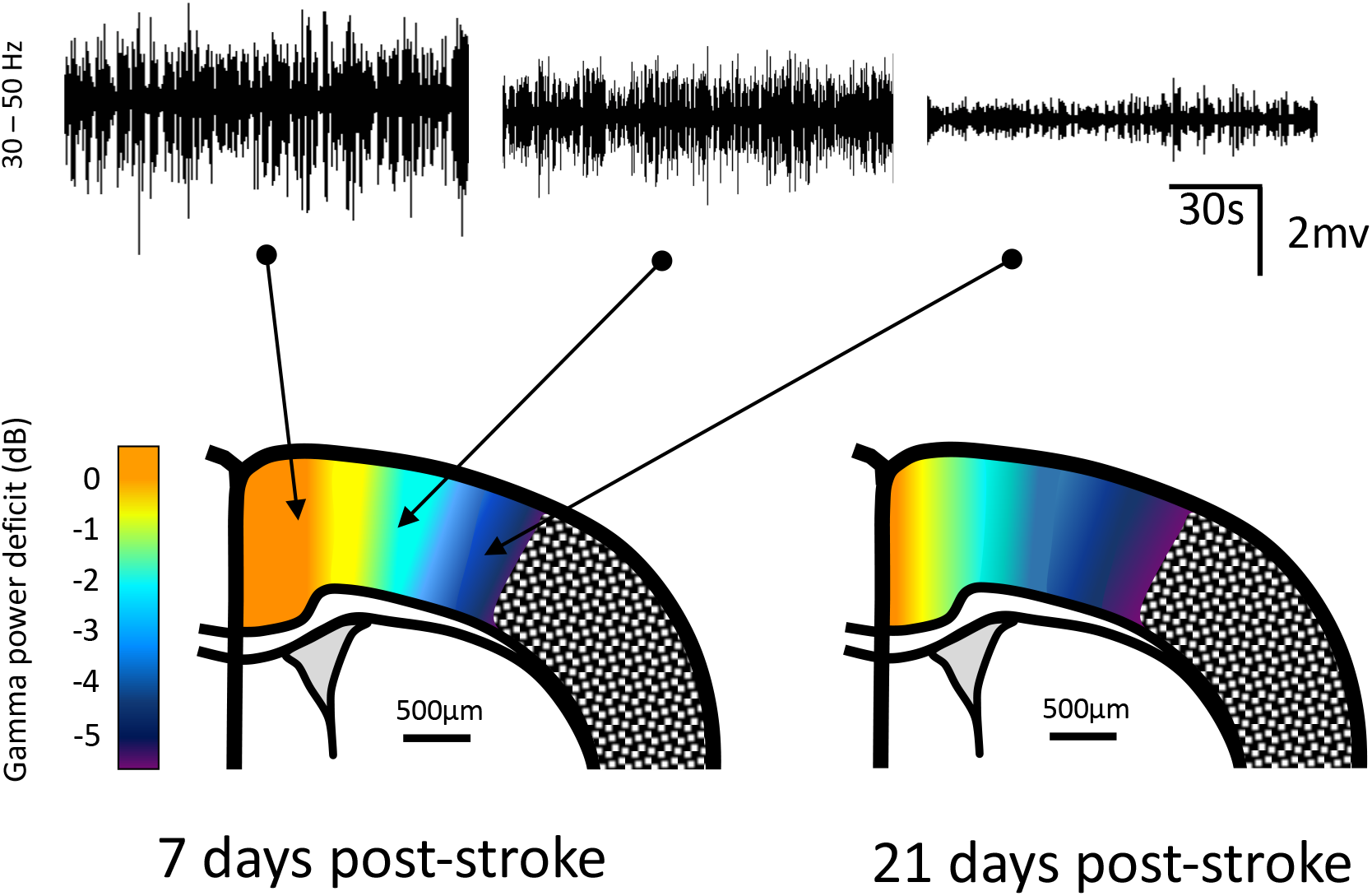
Model of gamma deficit in the cortical environment of an ischemic lesion. Based on the 2 power functions best describing the significant correlations obtained in Figure 3, a spatial modelization of gamma power deficit in the peri-infarcted cortex was drawn. Top: three examples of raw LFP band-pass filtered (30-50Hz) recorded at various distances from the infarction, 7 days after MCAO.

## Discussion

The main finding of this study is that the cortex surrounding an ischemic lesion undergoes a long-lasting deficit in gamma (30-50 Hz) oscillatory power, the latter being an emerging neurophysiological determinant of neural communication and plasticity.

In human, the highest potential for recovery is observed within the first 3 months after stroke (Biernaskie et al., 2004). In rodent models of focal cerebral ischemia, this critical period of plasticity corresponds to the first 4 weeks, after which functional recovery gains are no more significant (Brown et al., 2007; Dijkhuizen et al., 2001).

We chose to study the spontaneous oscillatory activity of the repairing cortex under anesthesia to avoid the stochastic changes of the oscillatory pattern associated with the awake state and favor the stability and the reproducibility of an internally driven regimen (Lustig et al., 2016). Isoflurane is a well-known depressor of cortical activity (Land et al., 2012), however at ~1%, we systematically observed a LFP oscillation pattern composed of three peaks reminiscent of the delta, theta and gamma frequencies in the awake mice.

We found no clear differences with control animals for delta and theta power at 7 and 21 days post-stroke in the neighborhood of the lesion. Neither the necrotic environment nor the loss of proximal afferent fibers seems to prevent peri-infarct neural networks to oscillate at these frequencies. A similar conclusion could have been drawn for gamma oscillations when averaging their power over the entire peri-infarcted zone. However, our detailed analysis revealed an accelerating collapse of gamma power the closer we get from the lesion. The extent of the deficit spanned 2 mm away from the glial scar, a distance matching many previous measurement of the peri-infarct cortex in the rodent brain (Brown et al., 2007). In the human brain, the extent is not proportionally bigger, but it represents an important volume of grey matter with high plasticity potential (Funck et al., 2016). As for the relevance of 2 mm of cortex, it represents the equivalent distance of 5 to 6 cortical columns in mice (Tischbirek et al., 2019). Moreover, cortical motor microcircuits are functional units organized in territory of 100 μm receiving local inputs from ~500 μm away (Amirikian and Georgopoulos 2003). Of note, plastic reorganization of neurons starts readily after the edge of the glial scar, that is, in the present case, ~160 μm from the necrotic core (Brown et al., 2007). These topological considerations emphasize the potential importance of this 2 mm strip of cortex for post-stroke functional recovery.

What is the origin of the selective depression of cortical gamma rhythm around the consolidated lesion? Seven and 21 days after stroke, peri-infarct neurons are still under the influence of inflammatory processes and glial reactivity. It could be argued that cell swelling or water content in the tissue changes its conductance in a magnitude such as gamma oscillations are less detected. This is however unlikely as several studies verified that the cortical tissue conductivity tested with an intracortical electrode show weak frequency dependence similar to cerebrospinal fluid (Logothetis and Wandell, 2004; Miceli et al., 2017). Moreover, as suggested by Miceli et al. (2018) the filtering properties of the tissue itself should be a minor factor in shaping the LFP. Therefore, the origin of gamma depression in the periphery of the stroke lesion should be searched in the light of cell and neural network properties. Gamma oscillations are generated sporadically by local groups of cortical neurons spatially confined into the volume of a putative column (Steriade et al., 1996) that is approximately 250 μm in diameter (Tischbirek et al., 2019) beyond which the amplitude of a gamma oscillation decreases rapidly with distance (Sirota et al., 2008). Possibly, and in agreement with several studies suggesting patchy selective neuronal death into the surviving cortex, long after the consolidation of the ischemic lesion, the number of neurons required to generate enough gamma power is not met in this zone (Baron, 2005; Guadagno et al., 2008).

Two other considerations can serve as food for thought regarding the origin of gamma deficit. Firstly, gamma oscillations are known to emerge from the interaction of parvalbumin interneurons and principal cells (Buzsáki and Wang, 2012). A selective reduction of these interneurons in the ipsilateral perilesional cortex has been associated with an improved plasticity and training-induced functional recovery (Alia et al., 2016; Zeiler et al., 2013). Future studies on the specific survival of these interneurons (Baron, 2005) and the integrity of their perineuronal net (Härtig et al., 2017), will help to interpret the present observation. Secondly, several demonstrations have been made in the last decade showing that the hypoexcitability of peri-lesional neurons can be corrected by antagonists of GABAAR tonic inhibition to boost functional recovery (Clarkson et al., 2010). Whether this excess of tonic inhibition can dampen the power of gamma oscillations is not clear (ref?). Outside of the cortex at least, reducing tonic inhibition leads to the emergence of gamma oscillations in hippocampal CA3 slices (ref?).

Little is known about gamma oscillation changes after stroke, since most studies have been done using EEG or EcoG with low spatial resolution and poor detection of frequencies above 30 Hz (Lake et al., 2016; Rabiller et al., 2015). Here we found that gamma power is strongly and persistently depressed in a cortical area that closely matches with the peri-infarct zone described as a primary reservoir of neuronal plasticity. Since gamma abnormalities have been associated with many brain disorders of very different pathophysiologic background (Woo et al., 2010), it is tempting to consider that it is simply an epiphenomenon resulting from perturbation of cerebral cortical network functions, only useful as a good sensitive clinical readout (Woo et al., 2010). However, the role of gamma oscillations in synaptic plasticity is increasingly suggested. Synaptic strengthening or weakening required to build new neuronal networks as a result of learning –and most likely of *re-learning* after a brain lesion– is an activity-dependent process driven by the principle of temporal coincidence of pre- and post-synaptic firing (Hebb, 1949). Oscillation of neuronal membrane potential at the gamma frequency allows synchronized neuronal firing within a temporal window that is optimal for STDP process (Engel et al., 1992; König et al., 1996; Magee and Johnston, 1997; Bi and Poo, 1998; Buzsáki and Draguhn, 2004; Wespatat et al., 2004; Harris et al., 2003, 2005). In other words, under a gamma rhythm of oscillation, the probability to associate pertinent information to assemble a neuronal circuit, increases. In support of the role of gamma oscillation in plasticity, several studies have demonstrated that the power of hippocampal gamma oscillation during learning predicts the quality of the recall (Sederberg et al., 2003, 2007; Headley and Paré, 2017). More compelling demonstrations are certainly necessary to prove a causal role of gamma oscillations in cortical synaptic plasticity and it remains to be shown that gamma deficit is actually associated with poor plasticity and functional recovery. But based on this concept, a lot of efforts are being carried out to stimulate the human brain with transcranial alternative electric current (tACS) at a specific frequency supposed to entrain natural oscillations and boost network plasticity (see Bland and Sale, 2019 for a recent update). While many challenges are still ahead to understand the neurobiology behind the observed effect, encouraging data suggest that tACS delivered at gamma frequency can be used to improve motor performance in humans (Guerra et al., 2018).

The cease of inputs originating from the necrotic cortex causes a functional imbalance in many regions of the brain, not necessarily directly connected to dead neurons. This phenomenon named diaschisis, describes changes of neuronal activity that will positively or negatively contribute to the reorganization of brain circuits (Reggia, 2004). One of the most obvious expressions of diaschisis after a cortical stroke is the activity change of the homotopic contralateral area. Both clinical and pre-clinical studies have shown hyperactivity in the contralateral cortex due to disinhibition (Andrews, 1991; Brown et al., 2009; Dijkhuizen et al., 2001; Tombari et al., 2004; Cramer et al., 2011). However, this increase in “activity”, whether measured by an increase of firing rate, excitability or neurovascular signaling is not necessarily accompanied by an increase of neuronal synchronization and therefore of oscillatory power. In the present study, we found an increase of the broadband LFP power of the contralateral cortex, mostly 3 weeks after stroke. The origin of this power increase is unlikely due to a genuine enhancement of the activity of each frequency-band oscillator but most likely due to an enlargement of the neuronal population entrained into these rhythms, because 1/ while the homotopic contralateral cortex displays an increased power in delta and theta wavebands, the peri-infarct cortex displays no change in delta and theta power at 21 days, irrespective to the distance with the lesion, suggesting that theta and delta subcortical oscillators deliver stable inputs; 2/ it is well known that a stroke induced transcallosal disinhibition is amplified by a downregulation of GABA receptor (Buchkremer-Ratzmann et al., 1996; Kokinovic and Medini, 2018) and a concomitant up regulation of NMDA receptor (Qü et al., 1998) in the contralesional cortex. Whether the increase of oscillatory power in the contralateral hemisphere is associated with favorable or unfavorable recovery, remains to be clarified (Kokinovic & Medini, 2018; Marshall et al., 2000; Silasi and Murphy, 2014).

In conclusion, the present work leads to the hypothesis that gamma deficit may slow down the process of synaptic reorganization occurring during the period of poststroke recovery. Accordingly, this result argues for the usefulness of therapeutic intervention aimed specifically at boosting gamma oscillations in order to enhance post-stroke functional recovery.

## Acknowledgment

This work was supported by INSERM (U1239), Rouen University, the Fondation pour la Recherche sur les AVC, the Normandy Region and the European Union (3R project). Europe gets involved in Normandy with European Regional Development Fund (ERDF).

## Notes

The authors have declared that no conflict of interest exists.

